# Application of an NMR/crystallography fragment screening platform for the assessment and rapid discovery of new HIV-CA binding fragments

**DOI:** 10.1101/2023.12.01.569544

**Authors:** Stuart Lang, Daniel A. Fletcher, Alain-Pierre Petit, Nicola Luise, Paul Fyfe, Fabio Zuccotto, David Porter, Anthony Hope, Fiona Bellany, Catrina Kerr, Claire J. Mackenzie, Paul G. Wyatt, David W. Gray

## Abstract

Identification and assessment of novel targets is essential to combat drug resistance in the treatment of HIV/AIDS. HIV Capsid (HIV-CA), the protein playing a major role in both the early and late stages of the viral life cycle, has emerged as an important target. We have applied an NMR fragment screening platform and identified molecules that bind to the **N**-terminal domain (NTD) of HIV-CA at a site close to the interface with the **C**-terminal domain (CTD). Using X-ray crystallography, we have been able to obtain crystal structures to identify the binding mode of these compounds. This allowed for rapid progression of the initial, weak binding, fragment starting points to compounds **37** and **38**, which have ^19^F-pK_i_ values of 5.3 and 5.4 respectively.

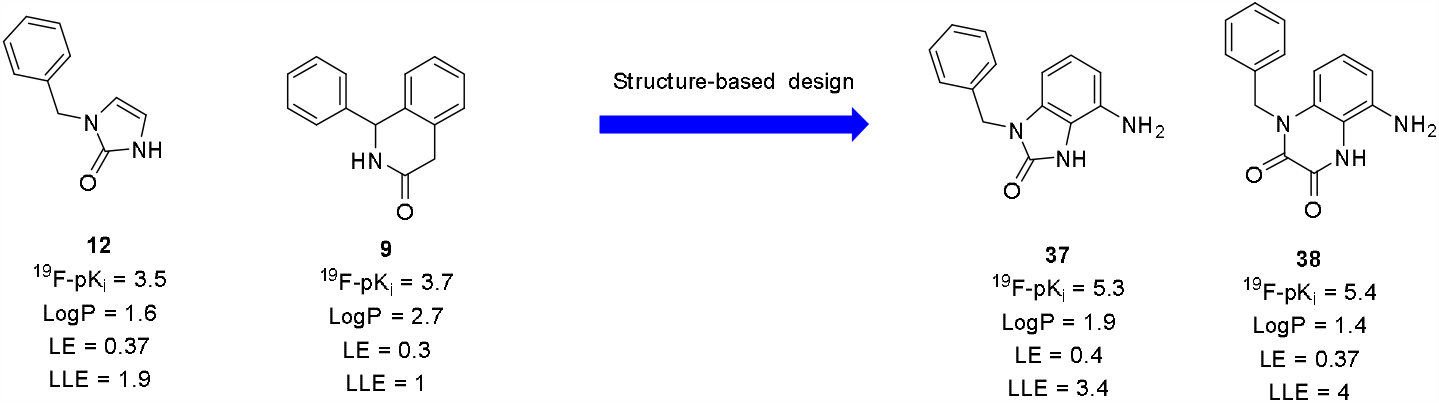

## Introduction

Acquired ImmunoDeficiency Syndrome (AIDS), first reported in 1981,^1^ is a major cause of human death. AIDS is caused by the Human Immunodeficiency virus (HIV), which is transmitted through sexual contact, exchange of body fluids (such as blood) or from an infected mother to her unborn child during pregnancy.^2^ In 2021 it was reported^3^ that there are 38 million people currently living with HIV, with 1.5 million new cases being reported annually, and 40 million people in total dying of AIDS/HIV related conditions: 0.65 million annually. HIV acts by mainly attacking the human immune system, infecting CD4^+^ T lymphocytes^4^ and reducing cell-mediated immune responses resulting in increased risk from other infections and diseases.^5^

Combination AntiRetroviral Therapy (cART) has been successfully used to control HIV in infected carriers, significantly reducing the viral load, resulting in an increased quality of life and increased life expectancy for patients. However, as HIV infection represents a chronic condition, long term use of cART can have disadvantages such as drug resistance, neurotoxicity, and liver failure.^6^ This means, despite many HIV infections currently being controlled by cART there is still an urgent need to develop alternative therapies that utilise novel strategies, including the identification of drugs that can interact with new targets or utilise unexplored mechanisms of action.

HIV-CA has attracted a lot of attention in recent years as a target for the treatment of HIV infections due to its role in various aspects of the viral life cycle.^7^ In the early stage of infection the CA protects the virus from the host’s immune system and mediates its entry into host cells. After entry into host cells the CA interacts with host factors such as Pin1, PDZD8, NUP153, cyclophilin A (CypA), cleavage and polyadenylation specific factor 6 (CPSF6), TRIM5a and MXB allowing for decortication of RNA and synthesis of complimentary DNA, beginning nuclear integration of the virus. These interactions are essential in allowing the virus to escape detection by the host’s immune system. This makes HIV-CA an attractive target for HIV replication, particularly in cases of drug resistant mutants.

HIV-CA has around 1500 copies of CA monomers, forming around 250 hexamers and exactly 12 pentamers, each being divided into two domains called the NTD and the CTD.^8^ These monomers are connected by interactions between the NTD and CTD along with interactions between the NTD and the NTD. Interactions between the CTD and CTD are responsible for the formation of the complete CA core.

There have been several inhibitors of HIV CA reported, with these being shown to bind to different sites within the CA structure (Figure 1). Ebselen (**1**),^9^ discovered using a time-resolved high energy high throughout screening (HTS-TR-FRET) assay, binds directly to the CTD and inhibits the early stages of the HIV life cycle. This molecule acts as a covalent inhibitor, binding to thiols in cysteines such as Cys198 and Cys218, forming a S-Se bond, giving way to ring opening and breaking of the Se-N bond.

**Figure 1.**
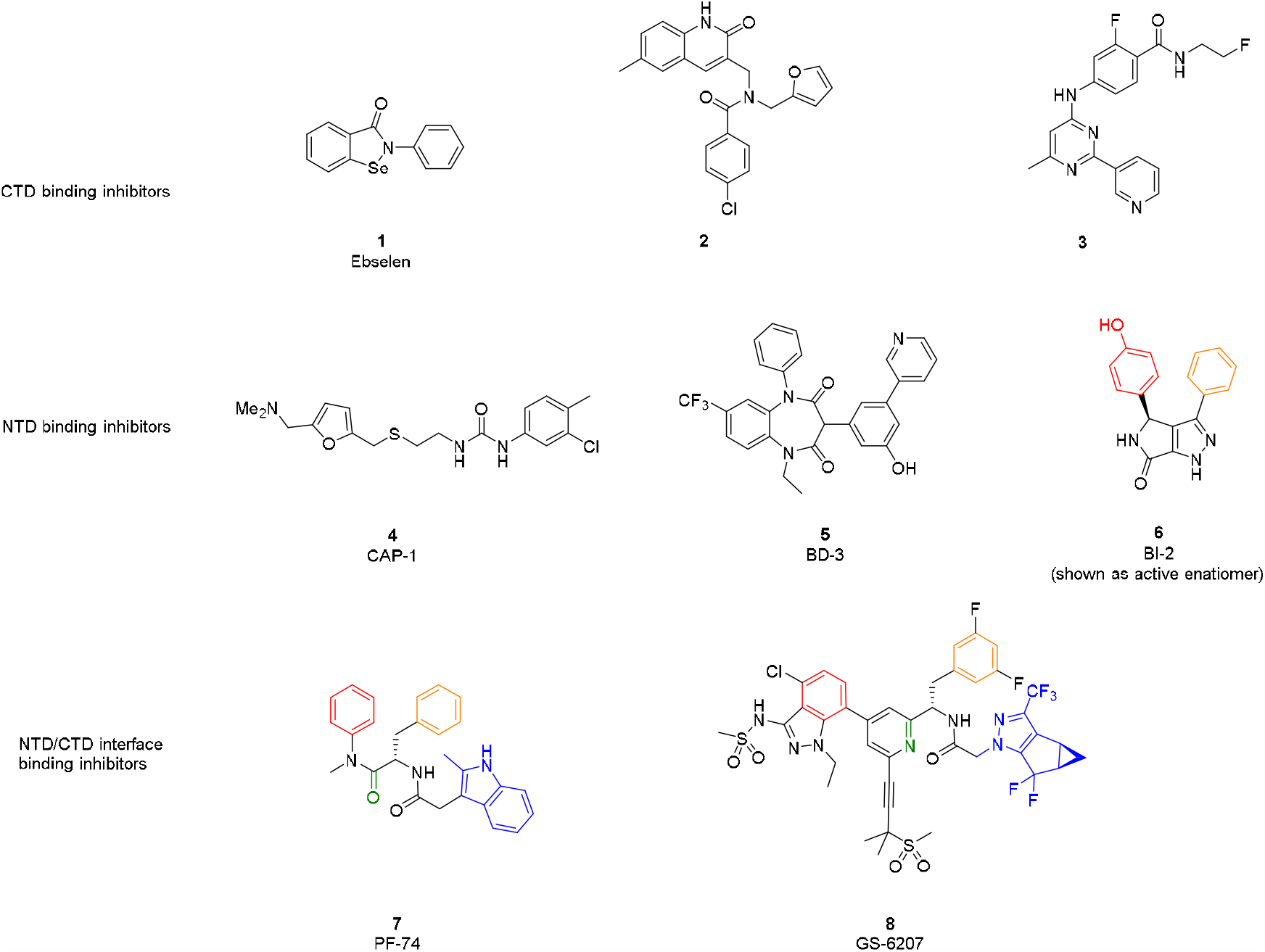
Public domain small molecules that are known to bind to HIV-CA.

Compound **2** was generated from a virtual screen^10^ that was carried out using structural information generated from NMR based techniques that studied the CA. Compound **2** has been shown to bind in the same region of the CA as CAI, a peptide inhibitor first reported in 2005.^11^ This region of the CTD spans helices 1 and 2 and has been shown to inhibit the assembly of mature CA. This CAI binding region was studied extensively, and compound **3** was generated as part of a bioisostere strategy, arising from hits identified as part of a HTS campaign.^12^ CAP-1 (**4**), also obtained through virtual screening, was the first HIV-CA inhibitor reported.^13^ In this case the virtual screen was carried out on the β-hairpin gap of the NTD. This compound works by disrupting the late stage of the virus’ life cycle, destroying polar interactions of the CA and inhibiting formation of the regular conical capsid. The aryl ring of **4** binds in a cavity formed by displacement of residue Phe32. The urea NH groups make an H-bond interaction with the carbonyl oxygen of main chain Val59 and the dimethylamino group interacts with the side chains of Glu28 and Glu29.

This pocket has been shown to be flexible, and subtly different conformations have been observed depending on the molecule located there. Many molecules, identified through HTS, bioisostere modifications and SAR explorations have been shown to bind to this pocket.^14^ One such compound, BD3 (**5**), has been shown to bind deep into the pocket with H-bonding interactions with the main chain NH on Phe32 and His62, along with water mediated H-bonding to the carbonyl group on main chain Val24 and Val59.^15^

From the screening of 60,000 compounds in a phenotypic cell-based assay BI-2 (**6**) along with its analogue BI-1 were identified.^16^ Through a high-resolution crystal structure, the binding site of this molecule was shown to be the same as that of the host factors CPSF-6 and NUP153. The phenol OH (coloured red in Figure 1.) of BI-2 makes a H-bond with Asn74, with the phenol ring. There are further H-bonds between the N-H of the lactam and Thr117, whilst the pyrazole ring makes an interaction with Asn57. This molecule binds to a site close to the NTD-CTD interface but is unable to make any interactions with the CTD. This molecule is unable to inhibit reverse transcription and is thought to inhibit viral replication by stabilising the CA..

PF-74 (**7**) can bridge across the NTD-CTD interface making interactions in both the NTD and CTD.^17^ Like BI-2 (**6**) it has two aryl rings (coloured red and orange in Figure 1.) that bind in the NTD and overlap with those coloured in BI-2; however, PF-74 has an indole system (coloured blue in Figure 1.) which is able to bind to the CTD, making an interaction with Arg173. PF-74 plays a vital role in disturbing both the early and late stages of the viral life cycle of HIV. Like BI-2 it can stabilise CA to trigger premature uncoating, but in addition to this, it is also able to inhibit reverse transcription, making the design and synthesis of molecules that are able to interact with both the NTD and CTD an attractive prospect.

PF-74 has been the most extensively studied HIV-CA inhibitor to date, with many groups creating analogues of this system.^18^ The most successful study resulted in the discovery of GS-6207 (**8**).^19^ This molecule, like PF-74, binds to the NTD along with the CTD. Steps have been taken to address the poor metabolic stability of PF-74 by replacing one of the labile amide bonds with a pyridine ring that acts as a bioisostere. Further groups have also been added to maximise lipophilic and H-bond donor/acceptor interactions, resulting in an increase in anti-viral activity. While the improved half-life of this molecule has meant that it only requires dosing once every 3 months, a complicated synthetic route leaves room for improvement with new HIV-CA inhibitors.

### Screening

Through the successful development of compounds that were able to interact at the NTD-CTD interface, as seen with PF-74 (**7**) and GS-6207 (**8**), and the inhibition that was observed at both early and late stages of the HIV life cycle we were keen to identify new molecules that could also bridge this interface. The HIV-CA protein was kindly provided by MRC Cambridge, using a method updated from Pornillos **et. Al**.20 We then screened molecules from the DDU fragment library (509 compounds) against HIV-CA using 3 different NMR methods; STD,^21^ WaterLOGSY^22^ and CPMG.^23^ For a compound to be progressed further, it had to show a positive binding event in the presence of HIV-CA, using all 3 NMR techniques, followed by a reduction of this event when PF-74 was added. This would ensure that not only was the compound binding to the HIV-CA,but that it was doing so at the same site as PF-74.

All putative hits gave a positive result in the STD experiment; however, this was not always confirmed in the other two NMR experiments (Table 1). A few examples of non-specific binding were observed: fragments displaying no change in peak intensity following the addition of PF-74. Most fragments were observed to bind competitively with PF-74 and were displaced following its addition.

**Table 1.** Overview of the DDU NMR fragment screen versus HIV-CA.

| Output | Number of Compounds | % of all compound |
| --- | --- | --- |
| Total compounds screened | 509 | - |
| Binding only observed in STD experiment | 17 | 3.3 |
| Binding in STD + 1 other experiment | 22 | 4.3 |
| Binding in all 3 experiments (STD, WaterLOGSY and CPMG) | 17 | 3.3 |

## Discussion

17 Compounds displayed binding in all 3 experiments (Figure 2) and were taken forward for further investigation. Binding was measured using a CPMG experiment, employing a ^19^F analogue of PF-74 as a spy ligand.^24^ Due to the low potency of these hits, structural information was sought to formally assess the interactions made by each compound with the HIV-CA protein. Attempts were made to obtain crystal structures for each of the compounds **9-25**.

**Figure 2.**
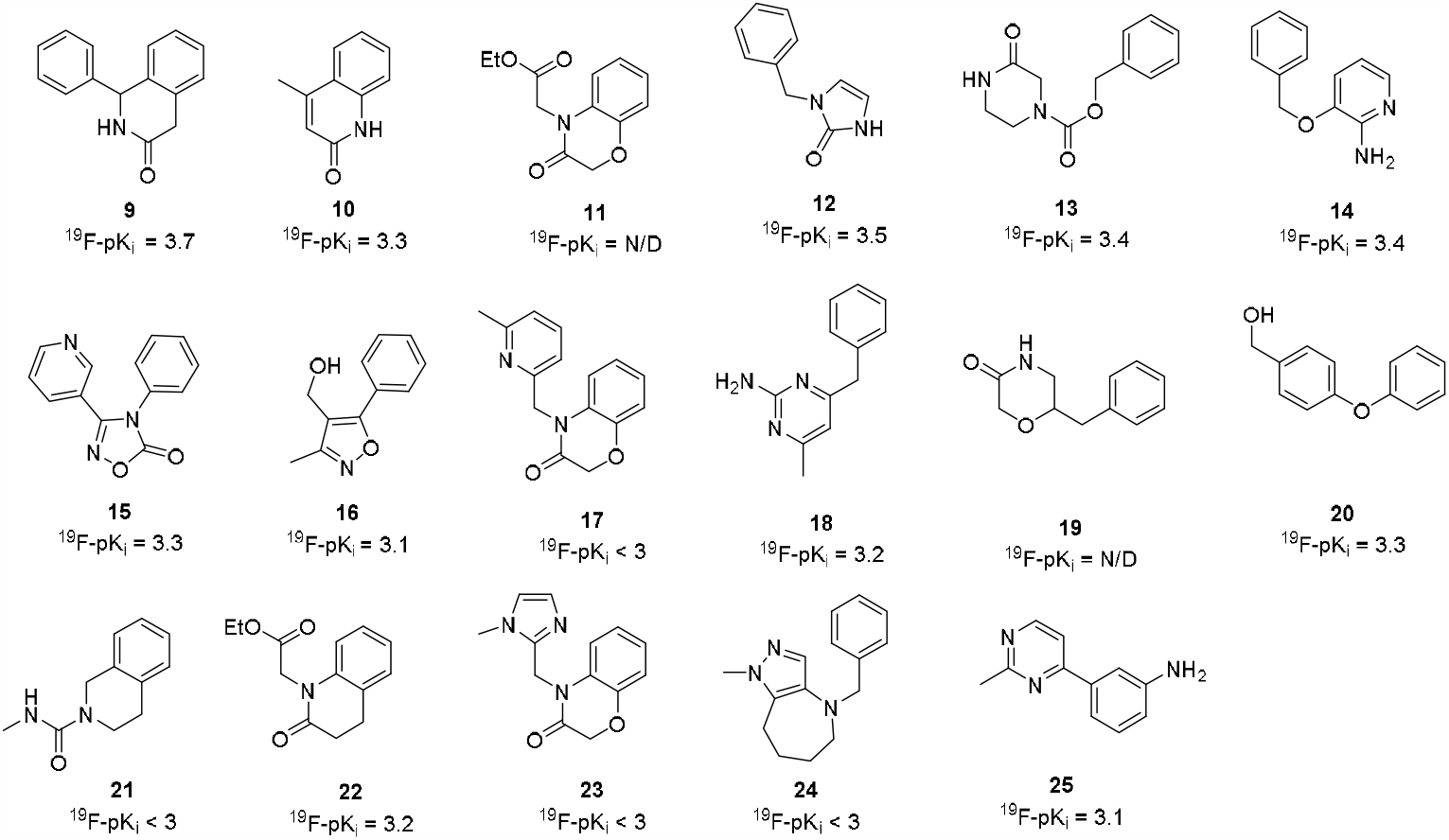
DDU Hit molecules that have shown binding in 3 NMR screening experiments: STD, WaterLOGSY and CPMG, as well as being competitive with PF-74.

Co-complex structures, generated by soaking of the HIV-CA crystals, were solved in space group P6 with resolutions of between 1.8 – 2.7 Å. All aligned closely with the CA:PF-74 crystal structure (rmsd of 0.3 – 0.7Å). The asymmetric unit is made of a single chain of CA, and the inter-protomer pocket is provided by a symmetry-related molecule. Gratifyingly, we were able to successfully obtain crystal structures for compounds **9-14** (Figure 3). All of these compounds bind to the same site as BI-2 at the NTD-CTD interface formed by α-helices 3, 4 and 5 of the NTD and the base of the CypA-binding loop from the CTD..

**Figure 3.**
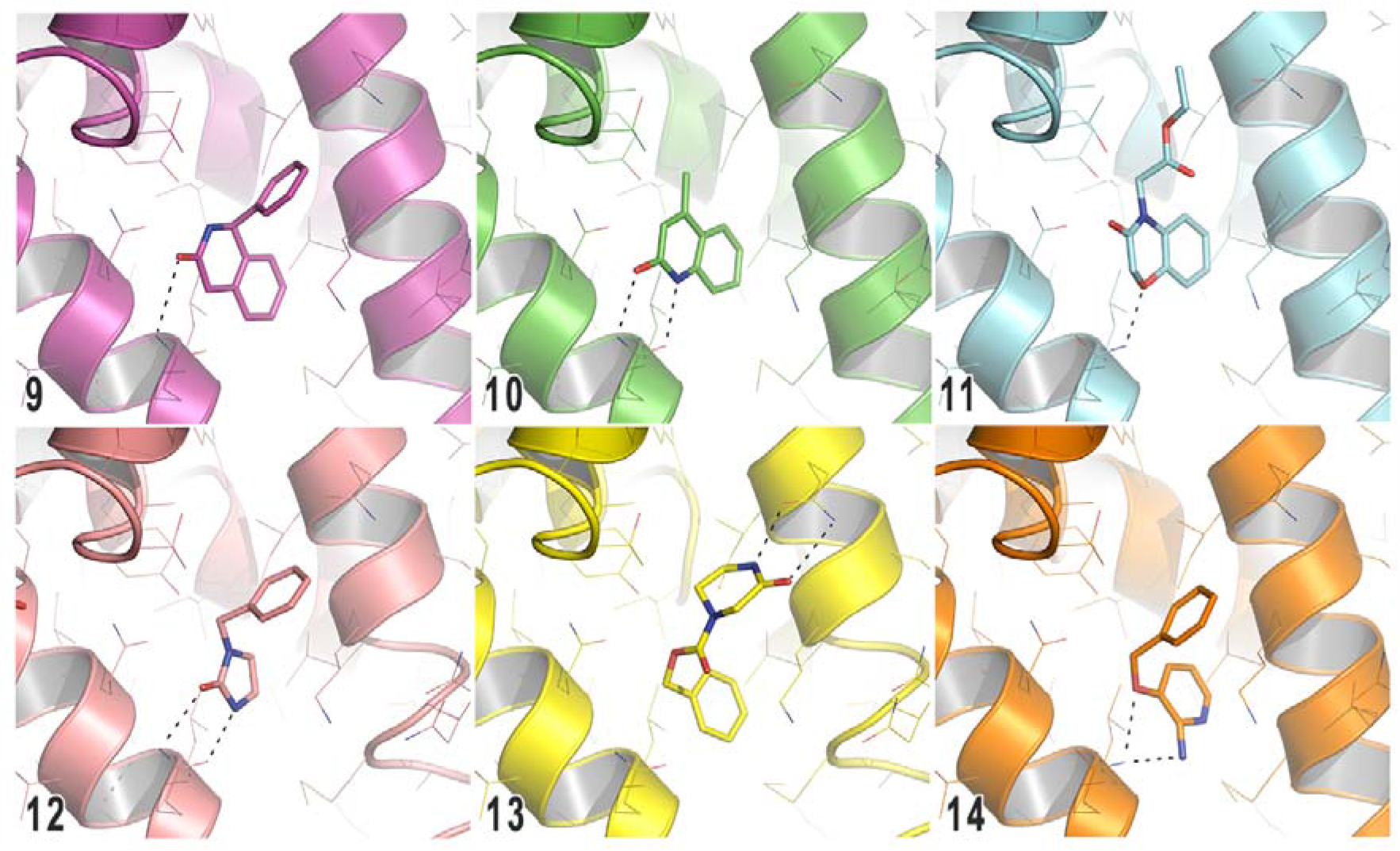
Crystal structure of compounds 9-14 in HIV-CA.

For Compound **9** (S-enantiomer), the binding mode is mainly hydrophobic, with only one weak H-bond (distance 3.4A) between the carbonyl and the side chain of the Asn57. Compound **10** also interacts with the side chain of Asn57 via 2 hydrogen bonds (2.7Å and 2.9Å). There is good overlay with the dihydroquinoline-3-one fused ring system of compound **9** and the quinolin-2-one system in compound **10**, with the carbonyl group making the same interaction with the CA protein in both instances. The same interaction to Asn57 is also observed with the imidazole-2-one group of compound **12**. Compound **13** can make a H-bond interaction via its lactam NHto Asn74. This is the same residue that made an interaction with the phenol OH in BI-2 (**6**). In addition to this, compound 13 can also make a water mediated H-bond interaction with Asn57 and a direct H-bond with Asn53 through the carbonyl of its carbamate.

The phenyl rings of compound **12**, and the phenyl ring of compound **9**, are situated in the hydrophobic region, where they can make interactions with the aliphatic side chains of Thr107, Tyr130 and Ile73. The phenyl ring of Compound **14** also makes the same hydrophobic interactions, whilst the exocyclic NH_2_ and ether O make H-bonding interactions with Asn57. The Lys70 residue can interact with the π-system of the fused aryl rings of compound **9, 10** and **11**. Compound **11** also makes a H-bond interaction with Asn57 through its cyclic O and appears to display a water mediated H-bond to Asn57 through its carbonyl group.

From our analysis of these crystal structures, we looked next to merge the fragments to improve potency, LE and LLE efficiency metrics (Figure 4). It was noted that compound **12** did not have an aryl ring occupying the same area of space in the protein as the equivalent group in compounds **9, 10** and **11**. Adding this aryl ring gave compound **26**. This modification gave rise to an increase in binding affinity (^19^F-pKi going from 3.5 to 4.6), but there was no increase in LE and LLE.

**Figure 4.**
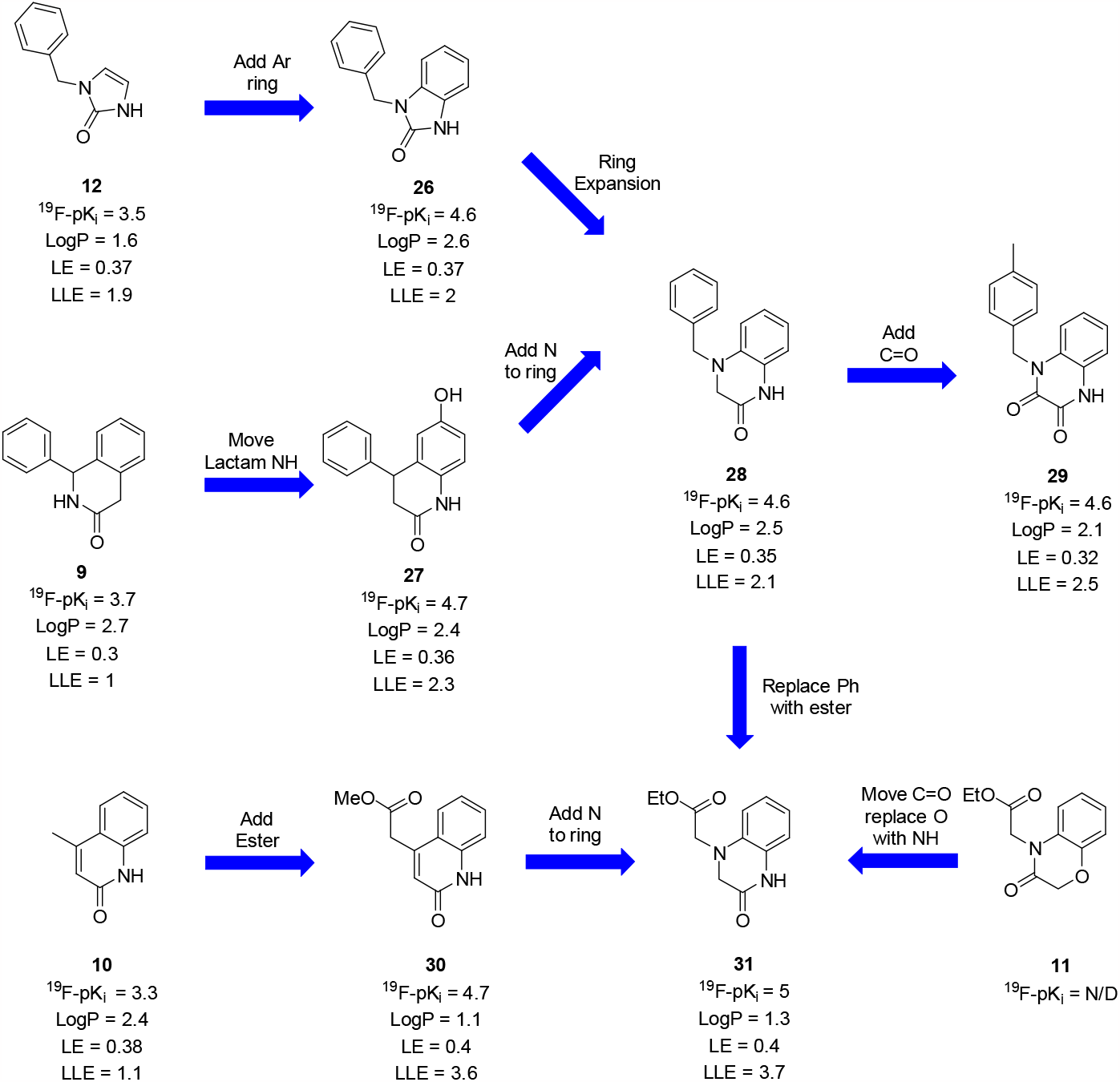
Optimisation of fragment cores

It was noticed in the crystal structure of compound **9** that only the carbonyl of the lactam was making a H-bond interaction with Asn57.. Moving the orientation of the carbonyl gave compound **27** which led to an increase in binding affinity from a ^19^F-pKi of 3.7 to 4.7. It was subsequently shown through further collection of crystal structures that compound **27** did not make additional H-bond interactions via its lactam NH (Figure 5). The observed increase in affinity can be explained by the additional H-bond between the -OH at position 7- and Asn74, causing the core to move into a different orientation.

**Figure 5.**
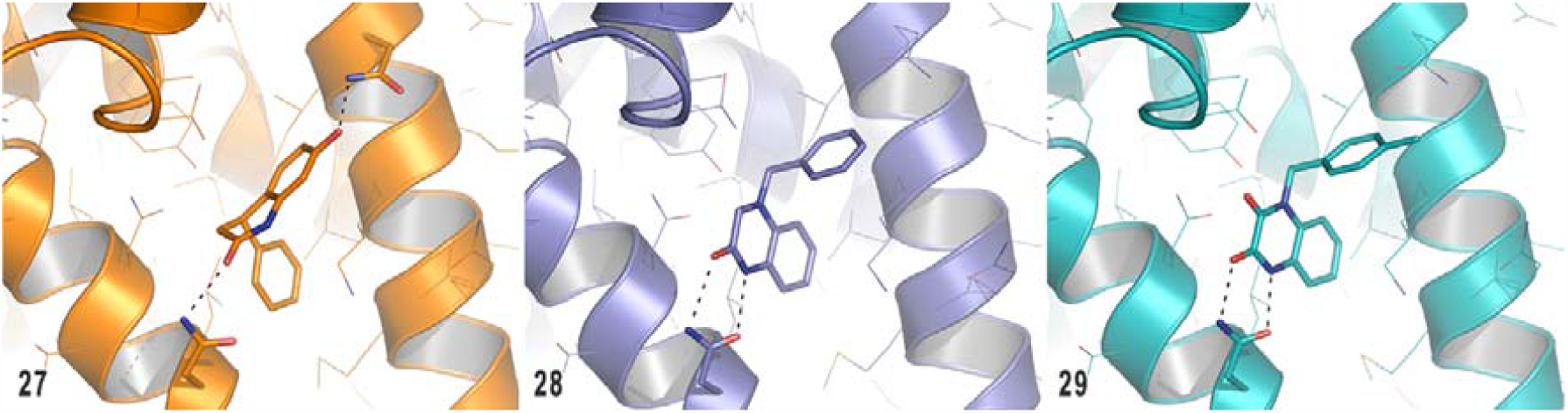
Crystal structure binding poses of compounds **27, 28** and **29**.

Compound **28** displayed a similar binding affinity, LE and LLE to compounds **26, 27**, and **29**. Compounds **28** and **29** bind in the expected orientation. The addition of an ester group to compound **10** gave compound **30** and resulted in an increase in binding affinity, whilst maintaining high ligand efficiency. Compound **31**, which has similarities to compounds **11, 28** and **30**, displayed a further increase in binding affinity (^19^F-pK_i_ = 5). The binding modes of compounds 30 and 31 were consistent (Figure 6) with the ester group overlaying with the aryl group in compounds **28** and **29** (Figure 5). The fused aryl systems of compounds **28, 29, 33**, and **37** were shown to overlay and occupy the same binding pose, with the cyclic NH and C=O groups orientated to interact with Asn57.

**Figure 6.**
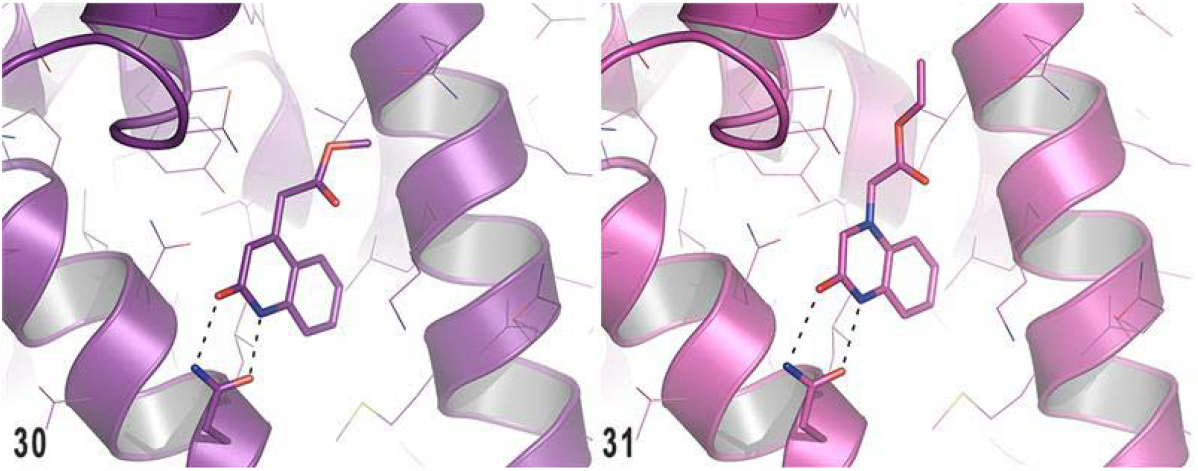
Crystal structure binding structure of Compounds 30 and 31.

We looked to further interrogate this section of the HIV-CA protein by performing a small scoping exercise, adding 1 or 2 atom groups to the 4- and 5-positions of the benzo-imidazol-2-one core, to assess the viability of growing the fragments from this position. The addition of a Br group at the 5-position (**32**, Table 2, entry 2) led to a small decrease in binding affinity, which coupled with the increase in lipophilicity and addition of a heavy atom resulted in a drop in LE and LLE. The addition of Br to the 4-position (**33**, Table 2, entry 3) saw an increase in binding affinity, however this change only maintained LE and LLE values comparable to the unsubstituted compound **26**.

**Table 2.**
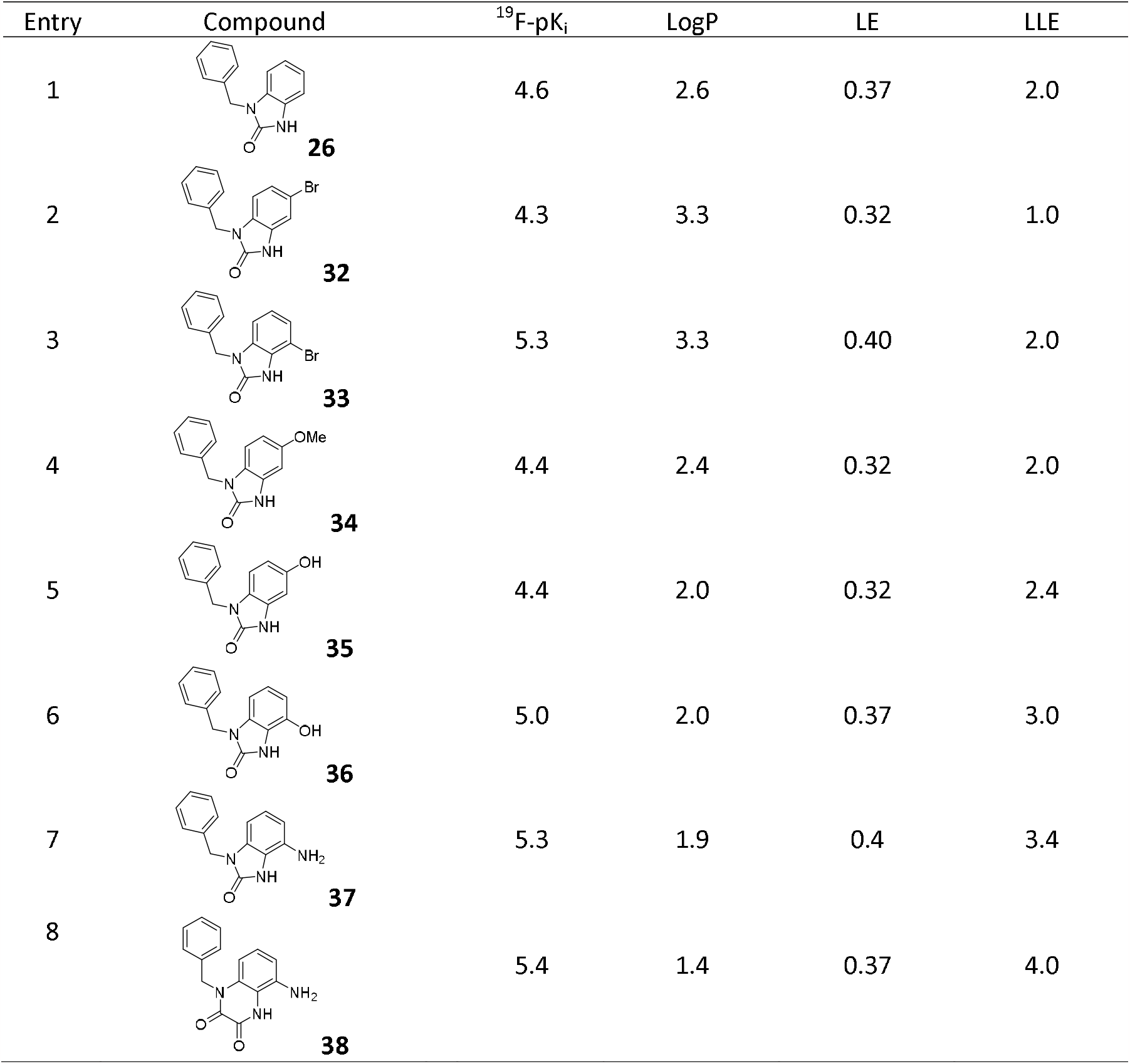
Addition of small groups to the periphery of fragment 26 at the 4- and 5- positions.

| Entry | Compound | <sup>19</sup>F-pK<sub>i</sub> | LogP | LE | LLE |
| --- | --- | --- | --- | --- | --- |
| 1 | **26** | 4.6 | 2.6 | 0.37 | 2.0 |
| 2 | **32** | 4.3 | 3.3 | 0.32 | 1.0 |
| 3 | **33** | 5.3 | 3.3 | 0.40 | 2.0 |
| 4 | **34** | 4.4 | 2.4 | 0.32 | 2.0 |
| 5 | **35** | 4.4 | 2.0 | 0.32 | 2.4 |
| 6 | **36** | 5.0 | 2.0 | 0.37 | 3.0 |
| 7 | **37** | 5.3 | 1.9 | 0.4 | 3.4 |
| 8 | **38** | 5.4 | 1.4 | 0.37 | 4.0 |

Further exploration of the 5-position with compounds **34** (-OMe) and **35** (-OH) saw no discernible change in binding affinity. However, the addition of an -OH at the 4-positon, as seen with compound **36**, resulted in an increase in binding affinity whilst maintaining LE comparable to compound **26**. The lowering of LogPcoupled with an increase in affinity, gave rise to an improved LLE of 3. Replacement of the OH group with an NH_2_ group gave compound **37**. Compound **37** demonstrated a further increase in binding affinity along with an increased LE and LLE. We then applied the knowledge gained from this series of compounds to prepare compound **38**, which has the same core as compound **29** (Figure 4).

Crystal structures were obtained for compounds **33** and **37** (Figure 7). Essentially, both molecules make the same H-bond interaction with Asn57 with hydrophobic interactions being made by the benzyl group with the aliphatic side chains of Thr107, Tyr130 and Ile73. The fused aryl rings of both molecules can make an interaction through their π–systems with the Lys70 residue. Despite the difference in nature of both groups at the 4-position (Br in **33** and NH_2_ in **37**), an increase in affinity (^19^F-pK_i_ change of +0.7 in both cases) is observed when compared to the unsubstituted compound **26**. Furthermore, these groups do no not appear to be making any significant interaction with the protein, but both display an improvement in LE.

**Figure 7.**
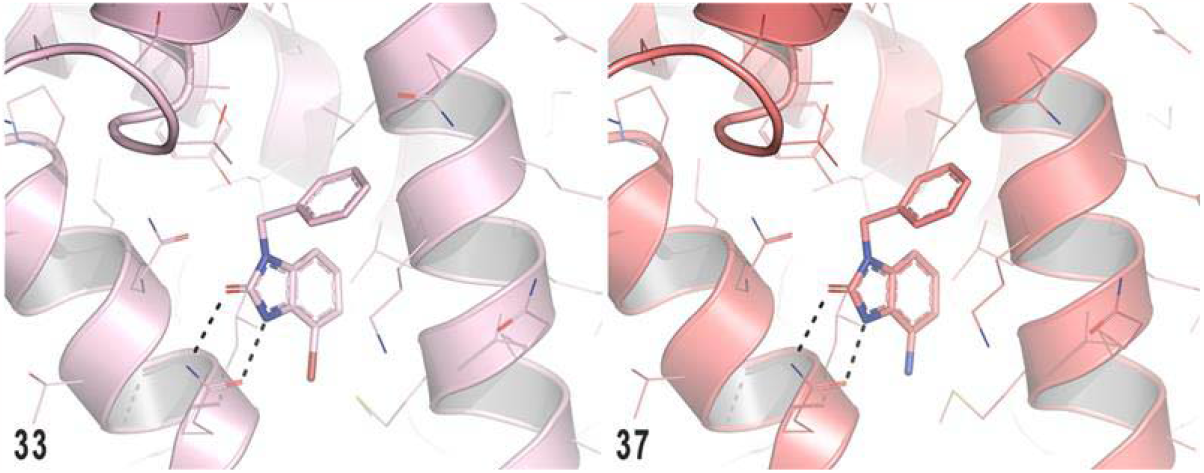
Crystal structure binding of Compounds 33 and 37.

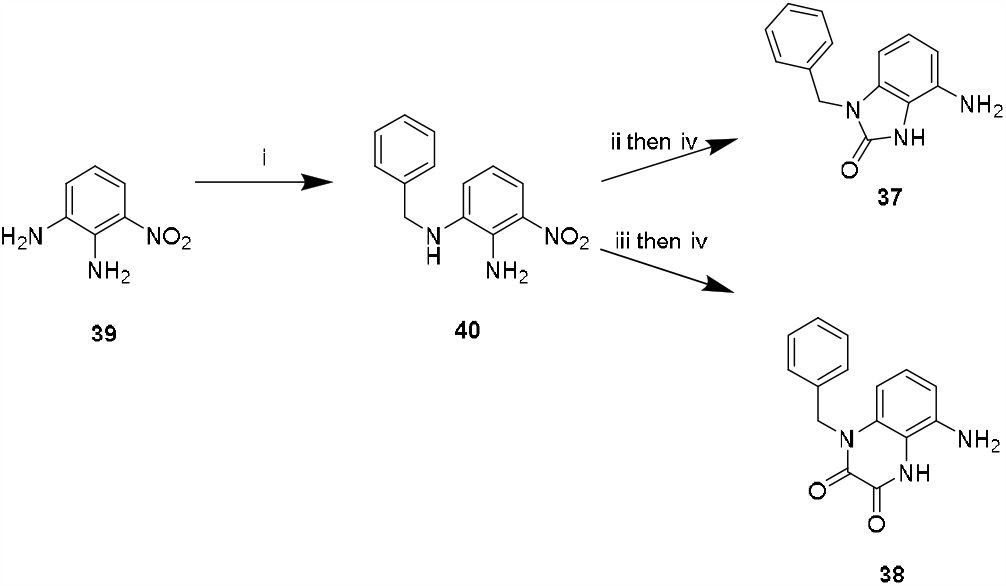
Reagents and conditions. (i) Benzyl bromide (1.2 eq), K_2_CO_3_ (1.5 eq), DMF, rt, 18h, 70%. (ii) CDI (1.6 eq), Et_3_N (2.5 eq), CH_2_Cl_2_, reflux, 4h. (iii) Oxalic acid (2 eq), AcOH/H_2_O (1:1), reflux, 5 days. (iv) NH_4_Cl (3-6 eq), Fe (3-6 eq), EtOH/H_2_O (4:1), reflux, 4h, 48% for **37** 27% for **38** (2 steps). **Scheme 1**. Preparation of Compounds **37** and **38**.

While compounds **9**-**36** are all commercially available and stored in the internal DDU compound collection, novel compounds **37** and **38** required synthetic preparation. This was readily achieved by addition of a benzyl group to 3-nitrobenzene 1,2-diamine **39** giving common intermediate **40** (Scheme 1). Compound **40** was subsequently treated with CDI followed by iron mediated reduction of the nitro yielding compound **37**. Alternatively, compound **40** can be treated with oxalic acid followed by iron reduction to give compound **38**.

## Conclusion

NMR based fragment screening has been successfully prosecuted against members of the DDU fragment library to identify new fragments binding to HIV-CA. Most compounds that gave a positive binding event in the experiments were displaced by PF74, indicating only low levels of non-specific binding to the HIV-CA protein. 17 of the 509 compounds screened were shown to bind to the protein in all three NMR experiments and were displaced by the inhibitor PF-74. This corresponds to a hit rate of 3.3%.

Following successful crystallisation of 6 compounds we were able to design new analogues and assess their binding. Using this information, structure-based design allowed rapid preparation of compounds **37** and **38**. Compounds **37** and **38** displayed improved ^19^F-pK i^’^s of 5.3 and 5.4 respectively. This process demonstrates that our NMR/crystallisation platform is not only capable of identifying and assessing weak binding, low molecular weight compounds, but it can also be applied to the design of novel compounds displaying improved binding affinities.

Due to the small size of compounds like **37** and **38** it is unsurprising that they are only capable of binding to the NTD of HIV-CA. However, it can be envisaged that this approach will allow compounds to progress from fragment-like to drug-like, binding not only to the NTD, but also with the CTD as has been observed with molecules like PF-74.

### Experimental

#### Protein Production

The HIV-1 capsid (HIV-CA) protein was kindly provided by MRC Cambridge, using a method updated from Pornillos **et. al**.^20^ The gene for HIV-1 capsid A14C/E45C/W184A/M185A (HIV-CA) was cloned into a pOPT vector with expression of HIV-CA under the control of the T7 promoter. The plasmid was transformed in **E. coli** C41 and the culture grown for 16 hours at 14 °C in 2XTY media with expression induced with the addition of IPTG to a final concentration of 0.4 mM. The cell was pelleted by centrifugation at 4200 g and then resuspended in 50 mM TRIS pH 8.0, 50 mM NaCl, 20 mM β-mercaptoethanol supplemented with protease inhibitors and bug buster. The cells were lysed by sonication and the insoluble fraction removed by centrifugation (40000 g, 1 hour, 4 °C). An ammonium sulfate (20% w/v) precipitation step was carried out on the soluble fraction, with 15-30 min the supernatant stirred constantly at 4 °C, the precipitated material was pelleted (40000g, 20 min, 4 °C). The pellet was resuspended in 100 mM citric acid pH 4.5, 20 mM β-mercaptoethanol and then dialysed against 100 mM citric acid pH 4.5, 20 mM β-mercaptoethanol with a minimum of 3 buffer changes. The complete HIV-CA hexamer is assembled through stepwise dialysis against 50 mM TRIS pH 8.0, 1M NaCl, 20 mM β-mercaptoethanol, then 50 mM TRIS pH 8.0, 1M NaCl and finally 50 mM TRIS pH 8.0, 40 mM NaCl. The HIV-CA hexamers were then purified by ion exchange chromatography (Hi-TRAP Q) pre-equilibrated with 50 mM TRIS pH 8.0, 40 mM NaCl and eluted on a NaCl gradient. Fractions of interest were pooled and then finally purified by gel permeation chromatography (Superdex S200 26/60) in 20 mM TRIS pH 8.0, 40 mM NaCl, where HIV-CA elutes at a volume consistent with a molecular weight of ∼150 kDa.

#### NMR Fragment Screen Assay

A library of 509 fragments were screened as pools of eight compounds per NMR tube at a concentration of 490 μM for each fragment. and were added to pH 7.4 45 mM potassium phosphate buffer and 45 mM, Deuterium Oxide (10 % per tube for NMR spectrometer stability) yielding final concentrations of, 2.1 μM **HIV-Capsid hexamer**. The tubes were submitted to the NMR spectrometer using the automation software ICON-NMR. Binding to the protein was determined using the Saturation Transfer Difference (STD),^21^ WaterLOGSY^22^ and CPMG (Carr, Purcell, Meiboom and Gill)^23^ experiments. Determination of specific binding was performed by re-running the experiments for each solution after the addition of 42.5 μM of the known inhibitor PF74 in DMSO-d_6_.

The fragments were screened as pools of eight compounds per NMR tube and were added to pH 7.4 potassium phosphate buffer, Deuterium Oxide (50 μL per tube for NMR spectrometer stability) and the protein solution to yield solutions with a final volume of 550 μL. This gave final concentrations of 490 mM for each fragment, 2.1 μM HIV-CA, 45 mM potassium phosphate and 45 mM KCl. All spectra were analysed by visual inspection and reprocessed where necessary using the Bruker software TopSpin v.3.2. The spectra for each pool of compounds were overlaid with the spectra of the individual samples within each pool (where available) to determine which compounds were binding to the protein and to measure changes in the peak intensities in the three spectra on addition of PF-74.

#### FAXS Assay

An ^19^F NMR based binding assay was developed following the work of Dalvit **et al**.^**24**^ on the FAXS (Fluorine chemical shift anisotropy and exchange for screening) method. In this experiment the ^19^F signal of a ‘Spy’ molecule, which binds to the protein of interest is examined. A ‘Control’ compound is also included which does not bind to the protein giving confidence that any changes in the Spy signal are due to a binding event and not changes in conditions such as variations in shimming efficiency. In this study the Spy ligand was designed based on a known weak inhibitor of the capsid hexamer, PF74 (see supporting information). The CF_3_ group was included due to the three-fold increase in signal intensity over a single fluorine which makes detection more straightforward.

Solutions of 25 μM Spy ligand and 25 μM Control with increasing concentrations (0 to 2.8μM) of protein in pH 7.4 45 mM potassium phosphate buffer and 45 mM, Deuterium Oxide (10 % per tube for NMR spectrometer stability) were prepared to produce a calibration curve for the change in peak intensity of the spy ligand peak with protein concentration. The protein concentration was then adjusted to approximately 2.2 μM to remove the flatter part of the curve and then various ligands were added at relevant concentrations to determine their ability to displace the Spy ligand and hence determine their K_I_ values (K_D_ for the spy ligand and its apparent binding constant K_D_ ^app^ based on its displacement by the competitor.

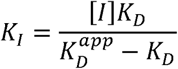

#### Crystallisation procedure

The first form of HIV-CA crystals are obtained using the sitting drop vapor diffusion method by mixing 200nl of HIV protein (12mg/ml) with 200 nl of mother liquor (0.1M Tris buffer, pH 8.0 to 9.0, 10-15% PEG550MME, 0.15M KSCN) with the latter as reservoir solution. These crystals were used to generate the structures by soaking crystals overnight in a 2ul drop of mother liquor supplemented with 1-10mM ligand (stock solution 20-200mM in 100 % DMSO).

Data were collected using a Rigaku M007HF copper-anode generator system equipped with Varimax Cu-VHF optics and a Saturn 944HG+ CCD detector or on Diamond Light Source (UK) synchrotron beamline. Data were processed using XDS^25^ and scaled with aimless from the CCP4 suite.^26^ Resolution limits were defined as CC half > 0.5.^27^ Structures were solved by molecular replacement using Phaser^28^ and the pdb entry 5HGM^29^ as the search model. Refinement was performed with iterative cycles of refinement using Phenix^30^ (or Refmac5^31^) and manual model building in Coot^32^ .

### Compound Preparation

#### General chemistry methods

Chemicals and solvents were purchased from commercial vendors and were used as received, unless otherwise stated. Dry solvents were purchased in Sure Seal bottles stored over molecular sieves. Unless otherwise stated herein reactions have not been optimized Analytical thin-layer chromatography (TLC) was performed on precoated TLC plates (Kieselgel 60 F254, BDH). Developed plates were air-dried and analysed under a UV lamp (UV 254/365 nm) and/or KMnO_4_ was used for visualization. Flash chromatography was performed using Combiflash Companion Rf (Teledyne ISCO) and prepacked silica gel columns purchased from Grace Davison Discovery Science or SiliCycle. Mass-directed preparative HPLC separations were performed using a Waters HPLC (2545 binary gradient pumps, 515 HPLC make-up pump, 2767 sample manager) connected to a Waters 2998 photodiode array and a Waters 3100 mass detector. Preparative HPLC separations were performed with a Gilson HPLC (321 pumps, 819 injection module, 215 liquid handler/injector) connected to a Gilson 155 UV/vis detector. On both instruments, HPLC chromatographic separations were conducted using Waters XBridge C18 columns, 19 mm × 100 mm, 5 μm particle size, using 0.1% ammonia in water (solvent A) and acetonitrile (solvent B) as mobile phase.1H NMR spectra were recorded on a Bruker Advance II 500 or 400 spectrometer operating at 500 and 400 MHz (unless otherwise stated) using CDCl_3_, DMSO-**d**_6_ or CD_3_OD solutions. Chemical shifts (δ) are expressed in ppm recorded using the residual solvent as the internal reference in all cases. Signal splitting patterns are described as singlet (s), doublet (d), triplet (t), multiplet (m), broadened (br) or a combination thereof. Coupling constants (**J**) are quoted to the nearest 0.1 Hertz (Hz). Low resolution electrospray (ES) mass spectra were recorded on a Bruker Daltonics MicrOTOF mass spectrometer run in positive mode. High resolution mass spectroscopy (HRMS) was performed using a Bruker Daltonics MicroTof mass spectrometer. LC−MS analysis and chromatographic separation were conducted with either a Bruker Daltonics MicrOTOF mass spectrometer connected to an Agilent diode array detector or a Thermo Dionex Ultimate 3000 RSLC system with diode array detector, the column used was a Waters XBridge column (50 mm × 2.1 mm, 3.5 μm particle size), and the compounds were eluted with a gradient of 5−95% acetonitrile/water + 0.1% ammonia, or with an Agilent Technologies 1200 series HPLC connected to an Agilent Technologies 6130 quadrupole LC/ MS, connected to an Agilent diode array detector, the column used was a Waters XBridge column (50 mm × 2.1 mm, 3.5 μm particle size) or a Waters X-select column ( 30 mm x 2.1mm, 2.5 μm particle size) with a gradient of 5-90% acetonitrile/water + 0.1% formic acid, or with an Advion Expression Mass Spectrometer connected to a Thermo Dionex Ultimate 3000 HPLC with diode array detector, the column used was Waters XBridge column (50 mm × 2.1 mm, 3.5 μm particle size) or a Waters X-select column ( 30 mm x 2.1mm, 2.5 μm particle size) with a gradient of 5-90% acetonitrile/water + 0.1% formic acid. All final compounds showed chemical purity of ≥95% as determined by the UV chromatogram (190−450 nm) obtained by LC−MS analysis.

### N-Benzyl-3-nitro-benzene-1,2-diamine (40)

3-nitrobenzene-1,2-diamine (5 g, 32.65 mmol) was dissolved in DMF (100 mL), Potassium carbonate (6.77 g, 48.98 mmol) and Benzyl bromide (6.7g, 39.18mmol) were added and the reaction mixture stirred at room temperature for 18 hours. The mixture was added to water (4 L) and extracted with CH_2_Cl_2_ (3 L). The solvent was removed, and the mixture was purified using silica chromatography, gradient elution using heptane to 50% EtOAc and Heptane, to give N1-benzyl-3-nitro-benzene-1,2-diamine **40**^**33**^ (5.57 g, 22.9mmol 70% yield) as a deep red powder.^1^H NMR (500 MHz, CDCl _3_): δ 7.71 (dd, **J**=1.2 and 8.7 Hz, 1H), 7.40-7.39 (m, 4H), 7.35-7.31 (m, 1H), 6.89 (d, **J**=7.6 Hz), 6.70 (dd, **J**=7.6 and 8.7 Hz, 1H), 5.98 (s, 2H), 4.33 (d, **J**=5.4 Hz, 2H), 3.47 (s, 1H); ^13^C NMR (125 MHz, CDCl_3_) δ 138.2, 137.8, 136.4, 133.4, 128.8, 127.9, 127.8, 117.7, 117.0, 116.6, 49.3; HRMS (ESI) calculated for C_13_H_14_N_3_O_2_ [M + H]^+^ 244.1081 found 244.1097; t_R_= 4.4 min.

### 7-Amino-3-benzyl-1H-benzimidazol-2-one (37)

3-Nitrobenzene-1,2-diamine (2.4 g, 9.87 mmol) was dissolved in CH_2_Cl_2_ (100 mL) and CDI (2.56 g, 15.79mmol) was added followed by Et_3_N (3.44 mL, 24.66 mmol). The reaction was allowed to stir at reflux for 18 hours. The mixture was diluted with in CH_2_Cl_2_ (100 mL) and washed with water (100 mL) and brine (100 mL). The mixture was purified by flash chromatography, gradient elution from heptane to 70% EtOAc/heptane mixture, to give 3-benzyl-7-nitro-1H-benzimidazol-2-one (2.29 g, 8.5 mmol, 86% yield) as a yellow solid, which was used directly in the next step. 3-Benzyl-7-nitro-1H-benzimidazol-2-one (2.29 g, 8.5 mmol) was dissolved in Ethanol (60 mL) and Water (15 mL) and Ammonium chloride (1.36 g, 25.51 mmol) and Iron (1.43 g, 25.51 mmol) added. The mixture was heated at reflux for 4 hours. The mixture was cooled and filtered through=h celite, then washed with EtOAc (250 mL) and washed with water (100 mL) to give **7-amino-3-benzyl-1H-benzimidazol-2-one** (1.13 g, 4.72 mmol, 56% yield) as a white solid. ^1^H NMR (500 MHz, DMSO-**d**_6_): δ 10.43 (1H, br s), 7.35 – 7.21 (5H, m), 6.68 (1H, t, **J** = 8.0 Hz), 6.32 – 6.26 (2H, m), 4.97 (2H, br s) and 4.92 (2H, s); ^13^C NMR (125 MHz, DMSO-**d**_6_) δ 154.5, 138.0, 132.3, 131.0, 128.9, 127.7, 121.9, 114.9, 107.9, 97.8 and 43.8; HRMS (ESI) calculated for C_14_ H _14_N_3_O[M + H]^+^ 240.1137 found 240.1135; t_R_ = 4.0 min

### 8-Amino-4-benzyl-1H-quinoxaline-2,3-dione (38)

Oxalic acid (4.07 g, 45.22 mmol) was added to a solution of N1-benzyl-3-nitro-benzene-1,2-diamine (5.5 g, 22.61mmol) in Water (65 mL) and Acetic acid (65mL). The mixture was heated at reflux for 5 days. The reaction was cooled to room temperature and the solvent removed by filtration. The mixture was recrystallized from CH_2_Cl_2_ (20mL) to give 4-benzyl-8-nitro-1H-quinoxaline-2,3-dione (2.73 g, 9.18 mmol, 41% yield) as a yellow powder. 4-Benzyl-8-nitro-1H-quinoxaline-2,3-dione (1 g, 3.36 mmol) was dissolved in Ethanol (30 mL) and Water (7.5 mL) and Ammonium chloride (1.08 g, 20.18 mmol) and Iron (1.13 g, 20.18 mmol) added. The mixture was heated at reflux for 4 hours. The mixture was cooled and filtered through celite, then washed with EtOAc (250 mL). Water (100 mL) was added. The precipitate was collected, and the water layer removed. The EtOAc layer was again with water (100 mL) and brine (100 mL). The solvent was removed, and the collected precipitates added to this to give **8-amino-4-benzyl-1H-quinoxaline-2**,**3-dione** (595 mg, 2.23 mmol, 66% yield) as a white powder. ^1^H NMR (500 MHz, DMSO-**d** 6): δ 11.20 (1H, br s), 7.35 – 7.19 (5H, m), 6.66 (1H, t, **J** = 8.0 Hz), 6.42 (1H, dd, **J** = 1.0 and 7.8 Hz), 6.35 (1H, dd, **J** = 1.0 and 8.0 Hz), 5.33 (2H, s) and 5.29 (2H, br s); ^13^C NMR (125 MHz, DMSO-**d**_6_) δ 157.2, 156.7, 138.6, 137.2, 129.0, 127.8, 127.4, 127.1, 122.2, 109.0, 103.5 and 46.0; HRMS (ESI) calculated for C_15_ H_14_ N_3_ O_2_ [M + H]^+^ 268.1086 found 268.1084; t_R_ = 3.8 min

