## Supplemental Table 1 for "Application of an NMR/crystallography fragment screening platform for the assessment and rapid discovery of new HIV-CA binding fragments"

| Complex | DDD00074110 | DDD00057456 | DDD00024969 | DDD00100452 | DDD00100439 | DDD00100555 | DDD01044153 |
| --- | --- | --- | --- | --- | --- | --- | --- |
| ID | 9 | 10 | 11 | 12 | 13 | 14 | 27 |
| PDB | 8QUB | 8QUH | 8QUI | 8QUJ | 8QUK | 8QUL | 8QUW |
| **Data measurement** | | | | | | | |
| Collection source | DLS-i03 | DLS-i03 | DLS-i03 | DLS-i03 | DLS-i03 | DLS-i03 | In house |
| Space group | P6 | P6 | P6 | P6 | P6 | P6 | P6 |
| Resolution range^a^ | 46.07-1.63  (1.66-1.63) | 56.82-1.55  (1.58-1.55) | 28.80-1.69  (1.73-1.69) | 56.80 - 1.63  (1.635-1.63) | 45.31 - 1.38  (1.40-1.38) | 56.68-1.67  (1.70-1.67) | 28.34-2.02  (2.13-2.02) |
| Unit cell dimensions | | | | | | | |
| a (Å) | 90.84 | 90.55 | 91.80 | 90.68 | 90.61 | 90.65 | 90.47 |
| b (Å) | 90.84 | 90.55 | 91.80 | 90.68 | 90.61 | 90.65 | 90.47 |
| c (Å) | 56.83 | 56.82 | 57.60 | 56.7 | 56.8 | 56.68 | 56.69 |
| α (°) | 90.00 | 90.00 | 90.00 | 90.00 | 90.00 | 90.00 | 90.00 |
| β (°) | 90.00 | 90.00 | 90.00 | 90.00 | 90.00 | 90.00 | 90.00 |
| γ (°) | 120.00 | 120.00 | 120.00 | 120.00 | 120.00 | 120.00 | 120.00 |
| Total observations | 327261 (13842) | 361157 (10669) | 286566 (14704) | 293642 (2882) | 409316 (4077) | 308200 (14100) | 99919 (9641) |
| Unique observations | 33459 (1676) | 38608 (1911) | 31130 (2247) | 30451 (334) | 51571 (1712) | 30894 (1526) | 17180 (2246) |
| Multiplicity | 9.8 (8.3) | 9.4 (5.6) | 9.2 (6.5) | 9.6 (8.6) | 7.9 (2.4) | 10.0 (9.2) | 5.8 (4.3) |
| Completeness (%) | 100.0 (100.0) | 99.9 (99.6) | 99.8 (98.6) | 91.3 (96.5) | 94.4 (63.2) | 100.0 (100.0) | 98.5 (89.7) |
| Mean I/sigma (I) | 19.2 (1.6) | 20.2 (1.3) | 20.4 (2.6) | 19.5 (2.0) | 24 (1.6) | 19.4 (1.8) | 11.0 (3.0) |
| R-merge (%) | 6.1 (129.8) | 5.5 (125.2) | 5.9 (62.3) | 6.3 (99.7) | 4.7 (64.1) | 7.0 (140.9) | 12.3 (47.9) |
| CC ½ | 99.9 (56.1) | 99.9 (52.0) | 99.9 (78.9) | 99.9 (69.9) | 99.8 (58.3) | 99.9 (56.1) | 99.7 (79.7) |
| **Refinement statistics** | | | | | | | |
| Resolution range | 46.07 – 1.63 | 56.82-1.55 | 28.80-1.69 | 56.80 - 1.63 | 45.31-1.38 | 56.68 – 1.67 | 28.34 – 2.02 |
| R-factor (R_work_/R_free_) | 18.9 / 21.0 | 17.8 / 19.5 | 17.3 / 21.6 | 17.7 / 19.9 | 14.2 / 17.9 | 17.5 / 20.9 | 18.9 / 23.0 |
| **Number of non-hydrogen atoms** | | | | | | | |
| Macromolecules | 1669 | 1667 | 1675 | 1833 | 1897 | 1672 | 1675 |
| Ligands | 17 | 12 | 17 | 13 | 17 | 15 | 18 |
| Solvent | 191 | 215 | 295 | 222 | 250 | 210 | 209 |
| RMS bond length deviation (Å) | 0.025 | 0.027 | 0.027 | 0.011 | 0.014 | 0.027 | 0.0181 |
| RMS bond angle deviation (°) | 2.07 | 2.26 | 2.12 | 1.72 | 1.71 | 2.27 | 1.769 |
| Ramachandran favored (%) | 97.6 | 97.2 | 98.1 | 97.9 | 98.3 | 97.7 | 98.1 |
| Ramachandran allowed (%) | 1.9 | 2.3 | 1.4 | 1.6 | 1.1 | 2.3 | 1.4 |
| Ramachandran outliers (%) | 0.5 | 0.5 | 0.5 | 0.5 | 0.6 | 0.0 | 0.5 |
| **Mean B-factor (Å^2^)** | | | | | | | |
| Macromolecules | 33.0 | 30.4 | 28.1 | 31.1 | 33.0 | 31.5 | 26.2 |
| Ligands | 38.9 | 36.6 | 47.9 | 52.7 | 44.4 | 39.8 | 30.7 |
| Solvent | 41.3 | 40.7 | 40.2 | 40.4 | 43.8 | 41.3 | 34.3 |

| Complex | DDD00100333 | DDD01728501 | DDD01728505 | DDD01728503 | DDD01829021 | DDD01829894 |
| --- | --- | --- | --- | --- | --- | --- |
| ID | 28 | 29 | 30 | 31 | 33 | 37 |
| PDB | 8QUX | 8QUY | 8QV1 | 8QV4 | 8QV9 | 8QVA |
| **Data measurement** | | | | | | |
| Collection source | In house | DLS-i03 | In house | In house | DLS-i04 | In house |
| Space group | P6 | P6 | P6 | P6 | P6 | P6 |
| Resolution range^a^ | 28.31 – 2.30  (2.42- 2.30) | 45.38 – 1.88  (1.91-1.88) | 45.17-2.20  (2.32 -2.20) | 28.33 – 2.70  (2.85 – 2.70) | 78-28 - 1.76  (1.81- 1.76) | 39.12 – 2.00  (2.05 – 2.00) |
| Unit cell dimensions | | | | | | |
| a (Å) | 90.39 | 90.76 | 90.35 | 90.40 | 90.39 | 90.34 |
| b (Å) | 90.39 | 90.76 | 90.35 | 90.40 | 90.39 | 90.34 |
| c (Å) | 56.62 | 57.00 | 56.58 | 56.65 | 56.41 | 56.42 |
| α (°) | 90.00 | 90.00 | 90.00 | 90.00 | 90.00 | 90.00 |
| β (°) | 90.00 | 90.00 | 90.00 | 90.00 | 90.00 | 90.00 |
| γ (°) | 120.00 | 120.00 | 120.00 | 120.00 | 120.00 | 120.00 |
| Total observations | 72763 (10425) | 117308 (4951) | 95928 (13980) | 42764 (6338) | 259387 (17402) | 56684 (4154) |
| Unique observations | 11849 (1717) | 21682 (1069) | 13511 (1977) | 7164 (1046) | 26170 (1924) | 17458 (1324) |
| Multiplicity | 6.1 (6.1) | 5.4 (4.6) | 7.1 (7.1) | 6.0 (6.1) | 9.9 (9.0) | 3.2 (3.1) |
| Completeness (%) | 99.9 (100.0) | 99.9 (98.1) | 100.0 (100.0) | 97.7 (98.8) | 100.0 (100.0) | 97.6 (99.3) |
| Mean I/sigma (I) | 7.1 (2.5) | 12.4 (1.6) | 5.5 (1.9) | 4.8 (1.9) | 11.6 (2.0) | 10.3 (3.0) |
| R-merge (%) | 22.1 (84.2) | 9.7 (103.4) | 29.2 (133.0) | 30.8 (100.6) | 10.8 (81.8) | 7.9 (42.4) |
| CC ½ | 98.9 (68.2) | 99.7 (52.7) | 98.7 (58.8) | 96.8 (67.6) | 99.9 (46.0) | 99.6 (69.5) |
| **Refinement statistics** | | | | | | |
| Resolution range | 28.31 – 2.30 | 45.38 – 1.88 | 35.33 – 2.20 | 28.33 – 2.70 | 50.23 – 1.76 | 39.12 – 2.00 |
| R-factor (R_work_/R_free_) | 20.1 / 27.5 | 17.9 / 21.5 | 22.2 / 28.7 | 23.0 / 30.2 | 20.2 / 24.5 | 20.5 / 25.4 |
| **Number of non-hydrogen atoms** | | | | | | |
| Macromolecules | 1696 | 1671 | 1704 | 1688 | 1694 | 1677 |
| Ligands | 22 | 20 | 16 | 17 | 18 | 18 |
| Solvent | 134 | 187 | 133 | 64 | 225 | 215 |
| RMS bond length deviation (Å) | 0.0069 | 0.0229 | 0.0083 | 0.0078 | 0.022 | 0.018 |
| RMS bond angle deviation (°) | 1.435 | 1.978 | 1.464 | 1.525 | 2.035 | 1.773 |
| Ramachandran favored (%) | 98.5 | 97.7 | 98.6 | 97.2 | 97.6 | 97.3 |
| Ramachandran allowed (%) | 0.5 | 1.8 | 1.4 | 2.8 | 2.0 | 2.2 |
| Ramachandran outliers (%) | 1.0 | 0.5 | 0.0 | 0.0 | 0.4 | 0.5 |
| **Mean B-factor (Å^2^)** | | | | | | |
| Macromolecules | 35.2 | 30.8 | 32.8 | 33.4 | 27.8 | 27.5 |
| Ligands | 37.2 | 48.3 | 43.4 | 23.8 | 26.7 | 34.1 |
| Solvent | 34.2 | 38.9 | 34.2 | 17.6 | 37.8 | 33.7 |

Table 1: Summary of data collection and structure refinement statistics. Values in

parenthesis are for the highest resolution shell. All measured data were included in

structure refinement.
